# JNK knockdown enhances CAR-T cell cytotoxicity

**DOI:** 10.1101/2025.08.14.666406

**Authors:** Charles J Kuhlmann, Chloe E Jepson, Madison T Blucas, Fatema Suleiman, Anusha Manda, Yoshiko N Kamata, Masakazu Kamata

**Affiliations:** Department of Microbiology, Heersink School of Medicine, University of Alabama at Birmingham, Alabama, USA; Department of Pathology, Department of Microbiology, Heersink School of Medicine, University of Alabama at Birmingham, Alabama, USA; O’Neal Comprehensive Cancer Center, University of Alabama at Birmingham, Birmingham, Alabama, USA

**Keywords:** JNK, NFATc1, CAR-T cell therapy, ovarian cancer, granzyme B

## Abstract

**Background:** Boosting the performance of chimeric antigen receptor T (CAR-T) cell therapy in solid tumors may provide a substantial advantage for cancer patients. Recognizing the vital role of the nuclear factor of activated T cells (NFAT) in T cell function, we hypothesized that the strategic regulation of NFAT activity by targeting c-Jun N-terminal Kinases (JNK) can bolster the tumor-eradicating potential of CAR-T cells.

**Methods:** We developed a lentivirally encoded short-hairpin RNA (shRNA) for stable knockdown of JNK in CAR-T cells. CAR-T cells targeting human epidermal growth factor receptor 2 (HER2) were produced from human peripheral blood. Functionality was tested *in vitro* and in two xenograft models of human ovarian cancer.

**Results:** JNK knockdown in CAR-T cells suppressed antigen-induced stimulation and helper T cell cytokine production, while enhancing anti-tumor cytotoxicity *in vitro* and in ovarian cancer xenograft experiments. Mechanistically, JNK knockdown led to elevated levels of granzyme B expression which could be recapitulated through overexpression of NFATc1, suggesting an NFATc1 dependent mechanism of action.

**Conclusions:** JNK signaling is a significant regulator of CAR-T cell cytotoxicity, offering a potential strategy to directly enhance CAR-T effectiveness in human cancer therapies.

## Background

Several advancements have been developed to expand chimeric antigen receptor T (CAR-T) cell efficacy into solid tumors. Using cytokine support^1^, combining with checkpoint blockade^2^, or transcriptionally modifying CAR-T cells^3–5^ is effective to boost CAR-T cell persistence, tumor infiltration, and stemness. Despite these initiatives, it remains unclear how to optimize the performance of CAR-T cells once they reach the tumor site, making reliable tumor control elusive.

Nuclear Factor of Activated T cells (NFAT) proteins feature a family of five distinct isoforms (NFATc1-c4 and TonEBP^6^) that may possess distinct functionalities. For example, NFATc1 is implicated in T cell cytotoxicity^7^, while NFATc2 promotes T cell exhaustion^8^. Thus, managing NFAT signal represents a target for enhancing CAR-T cell functionality. Current approaches dampen NFAT signaling to reduce excessive CAR-T cell activation^9^; however, precise methods to optimize NFAT signaling are still lacking.

The c-Jun N-terminal Kinases (JNK) are regulators of the NFAT pathway^10–12^. JNK negatively regulates NFATc1 by blocking its nuclear entry^10, 12^. Conversely, JNK enhances NFATc2 isoform transactivation by phosphorylating its N-terminal regulatory domain^11^. Thus, JNK signaling may contribute to diverse functional outcomes due to its interactions with NFAT isoforms. While JNK plays a vital role in T-cell priming^13^, its role in governing the functions of previously activated T cells, including CAR-T cells, remains limited.

We employed ovarian cancer xenografts as a model for optimizing CAR-T cell efficacy in solid tumors. Ovarian cancer is often diagnosed in a late stage with treatment resistance^14^, emphasizing the need for new salvage regimens. Ovarian cancers are heavily immune-infiltrated but typically do not respond to checkpoint blockade therapy, partially due to low rates of tumor-specific T cells^15^. CAR-T cell therapy overcomes this obstacle by providing tumor-targeting T cells, but current clinical response rates in ovarian cancer therapy are low (∼20%)^16^. Thus, the CAR-T cell approach can be practical for ovarian cancer but requires further optimization.

We hypothesize that regulating CAR-T cell NFAT signaling through JNK suppression enhances the functionality of CAR-T cells against ovarian cancer. In this regard, we performed an in-depth analysis of the effects of JNK signaling in CAR-T cells by exploring how JNK knockdown relates to their stimulation, function, and repercussions on anti-tumor efficacy *in vitro* and *in vivo* models of ovarian cancer.

## Materials and Methods

### Study Design

We assessed the impact of JNK suppression on CAR-T cell stimulation and anti-tumor efficacy using xenograft ovarian cancer models and human cell cultures. CAR-T cells were produced from peripheral blood T cells of healthy donors.

### Cell line culture

HEK293T, SKOV3, OVCAR8, and Jurkat cell lines were obtained from ATCC. All cell lines were maintained in Isocove’s Modification of Dulbecco’s Medium (IMDM) (Cytiva #SH30259.02) with 10% fetal bovine serum (Omega Scientific #FB-02), 1% glutamax (Gibco #35050-061), and 1% antibiotic-antimycotic solution (α/α, Omega Scientific #AA-40). Cultures were maintained at 37°C in a 5% CO₂ incubator.

### Lentiviral vectors

Lentiviral particles were generated in HEK293T cells via calcium phosphate transfection^17^, concentrated 200-fold by centrifugation, and stored at -80°C. CAR vectors were constructed as shown (see **Fig.1D**). Briefly, the vector co-expressed shRNAs (targeting JNK or scramble) under the H1 promoter, ZsGreen under the EF1α promoter, and CAR protein under the MSCV promoter. For NFATc1 overexpression, the vector encodes NFATc1-P2A-mCherry under the EF1α promoter. The lentiviral NFAT-Luciferase reporter (NFAT-Luc) vector includes nine repeats of antigen receptor response element-2 (ARRE-2)^18^ driving Akaluc^19^ expression with a minimal IL-2 promoter (see **Supplemental Fig.1A**).

**Fig. 1:**
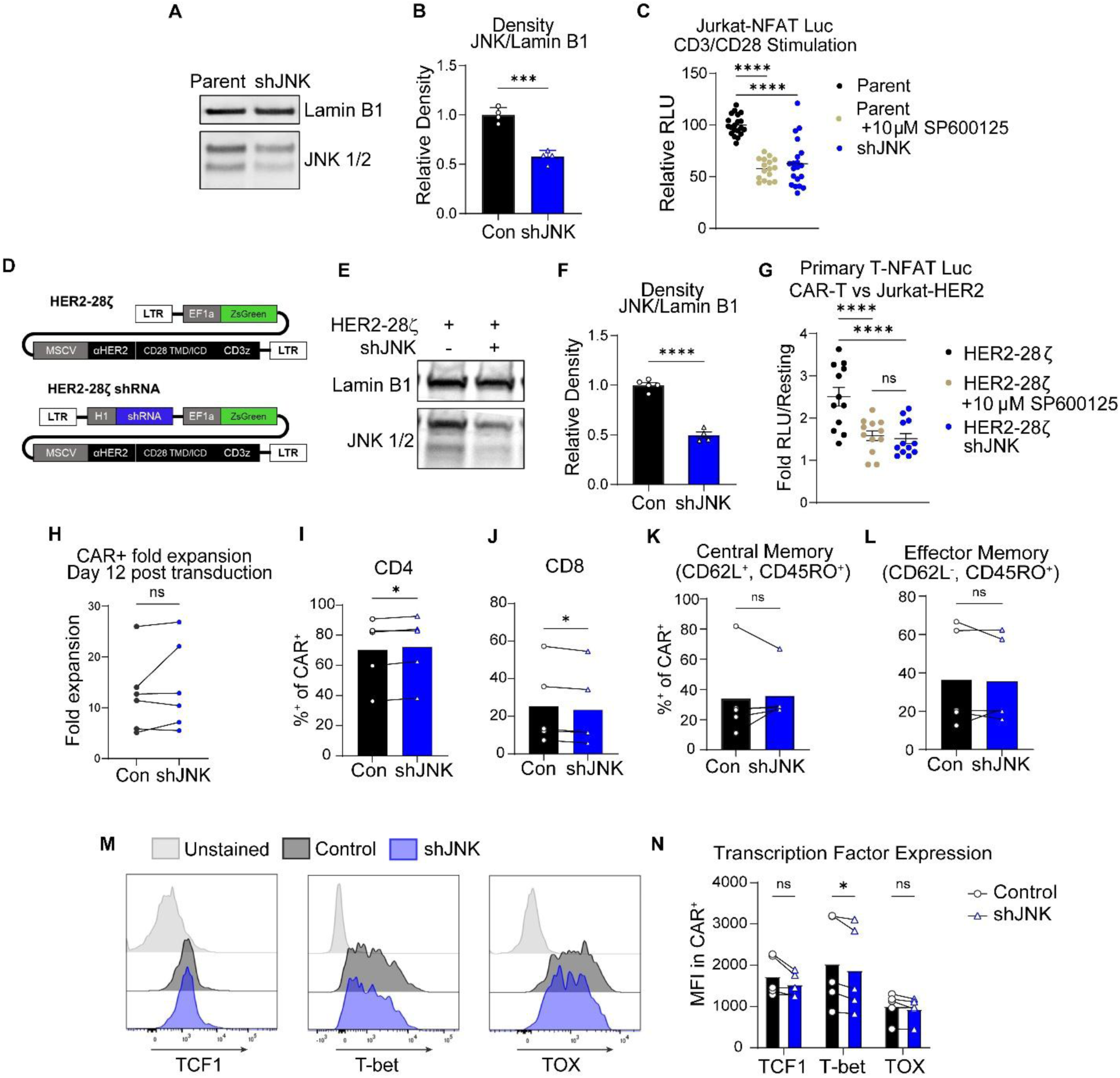
JNK knockdown suppresses CAR-induced NFAT signaling without impacting production. **A-B.** Jurkat NFAT-Luc cells were transduced with an shJNK encoding vector and JNK levels were analyzed by western blot. N = 2 experiments, 2 technical replicates each. Blot image (**A**) shows total JNK 1/2 and LaminB1, with densitometric quantification in (**B**). Unpaired T Test with Welch’s Correction. **C.** Luminescence assay of cells from (**A**) after stimulation with 2.5% w/v Immunocult CD3/CD28 antibody complexes +/- 10µM SP600125. N = 5 independent experiments. Aggregated data. Each data point shows percent of luminescence relative to the average of parental wells. One-way ANOVA with Fisher’s LSD test. **D.** Schematic of lentiviral vectors encoding CAR with or without shRNA cassette, which encodes either shScramble or shJNK. **E-F.** CAR-T cells were generated with control or shJNK vectors and analyzed by western blot on day 12 after transduction. N=2 donors, 2-3 technical replicates each. Aggregated data. Blot image (**E**) shows total JNK ½ and LaminB1, with densitometric quantification in (**F**). Unpaired *t-*test with Welch’s Correction. **G.** CAR-T cells were generated with control or shJNK vectors and co-transduction with NFAT-Luc vector. Luminescence assay after co-incubation with Jurkat-HER2 cells at a 1:1 ratio +/- 10µM SP600125. N=3 donors, 4 technical replicates each. Aggregated data, each data point shows fold luminescence over resting wells. One-way ANOVA with Fisher’s LSD test. **H.** Fold expansion of Control or shJNK CAR-T cells on day 12 post transduction. n=6 donors, aggregate data. Ratio paired *t-*test. **I-N.** CAR-T cells analyzed by flow cytometry on day 12 after transduction. N=5 donors. Ratio paired *t-*tests. Plots show the percentage of ZsGreen^+^ CAR-T cells expressing the indicated markers: CD4 (**I**), CD8 (**J**), CD62L^+^/CD45RO^+^ central memory cells (**K**), or CD62L^-^/CD45RO^+^ effector memory cells (**L**). Histogram plots (**M**) show intracellular staining of the indicated factors, with MFI quantification in (**N**).

### Human Peripheral Blood Mononuclear Cells (PBMCs)

Fresh leukocyte retention system (LRS) cones (LifeSouth Community Blood Centers, Gainesville, FL) were processed to obtain PBMCs by phosphate buffered saline (PBS) washing and red blood cell (RBC) lysis (PharmLyse, BD Biosciences # 555899). Cells were cryopreserved with Bambanker Freezing Media (Bulldog Bio # BB01).

### Primary T cell culture and assays

Cells were cultured in IMDM supplemented with 5% (v/v) Human AB Serum (Omega Scientific #HS-25), 1% glutamax, and 1% α/α. Activated T cells were generated by 36 h incubation of 6 × 10⁶ PBMCs with 80 µL anti-mouse IgG beads (Spherotech #MMSFC30-10) coated with 2µg αCD3 (Biolegend #300333) and 0.5µg αCD28 (Biolegend #302943) antibodies, followed by magnetic isolation. For lentiviral vector transduction, activated T cells were incubated for 2 hours with 50µL of concentrated lentiviral vector in 1mL medium per 1×10^6^ T cells in the presence of 8µg/mL polybrene (MilliporeSigma #H9268). T cells were expanded in medium containing 5ng/mL IL-15 and IL-7 for 12 days. CAR-T cell transduction was monitored by ZsGreen expression. Unless otherwise stated, all assays used 1.5 × 10^5^ T cells per well in 96-well plates with 25 IU/mL recombinant human IL-2 and stimulation as described in the figure legends. Stimulating agents include Immunocult αCD3/αCD28 antibody complexes (Stem Cell Technologies #10971), Biotinylated Recombinant Human HER2-Fc Chimera (Biolegend #796506), αCD3 antibody, and αCD28 antibody. JNK inhibitor SP600125 (MedChem Express # HY-12041) was used as indicated. Experiments on blood derived T cells without activation (see **Supplemental figure 3**) used T cells purified from thawed PBMCs with EasySep Human T-cell Enrichment Kit (Stem Cell Technologies #19051).

### Cytotoxicity assay

Firefly luciferase labeled^20^ SKOV3 and OVCAR8 cell lines were plated at 1×10^4^ cells/well, and co-cultured with flow-sorted CAR^+^- or untransduced T cells at the indicated effector:target (E:T) ratio. After 24 hours, D-Luciferin was added, and luminescence was measured with a Varioskan Lux spectrophotometer (Thermoscientific #VL0L00D0). Specific killing was calculated as: (1-(Luminescence in CAR-T well/average luminescence in Mock T cell wells for that E:T)) x 100. Assays using purified CD4^+^- or CD8^+^-CAR-T cells compared CAR-T containing wells to wells without T cells to calculate total killing.

### Flow cytometry

Antibodies used are listed in Supplementary Table 1. Zombie UV (Biolegend #423108) or Zombie Aqua (#423102) dye was used according to the manufacturer’s instructions to discriminate live and dead cells. Antibody staining was performed according to the manufacturer’s instructions in PBS supplemented with 2% FBS, 0.3% sodium azide, and 0.5 mM ethylenediaminetetraacetic acid. True-Nuclear Transcription Factor Buffer Set (Biolegend #424401) was used to enable transcription factor staining. Granzymes were stained using Thermo FIX & Perm (Thermo Fisher #GAS004) according to the manufacturer’s instructions. Monensin (Biolegend #420701) was used to enable surface CD107α staining. Samples were fixed in 2% paraformaldehyde (PFA) in PBS before acquisition. Data was acquired on BD LSR FORTESSA or BD FACS SYMPHONY A5. Data was analyzed using FlowJo 10 (FlowJo, LLC, OR). Flow sorting was performed using a BD FACS ARIA instrument.

### Mouse husbandry

The animal research described in the study was approved by UAB Institutional Animal Care and Use Committee (IACUC) and conducted following guidelines for the housing and care of laboratory animals of the National Institutes of Health and Association for Assessment and Accreditation of Laboratory Animal Care International. The mice used in this study are NOD.Cg-*Prkdc^scid^ Il2rg^tm1Wjl^*/SzJ (NSG) mice between 8 and 10 weeks of age. All animals were housed in autoclaved cages with isolators with individual air sources, supplemented with irradiated standard rodent chow (LabDiet, 5LJ5) and Hydropac.

### *In vivo* efficacy assay using human ovarian cancer xenograft mouse models

For SKOV3, NSG mice were engrafted with 1×10⁶ mCherry^+^-, firefly luciferase^+^-SKOV3 cells intraperitoneally (IP). After 14 days, 1×10⁶ CAR^+^-T cells were injected intravenously (IV). Tumor burden was measured weekly by bioluminescent intensity (BLI) after intraperitoneal injection of D-Luciferin and imaging (IVIS Lumina III, Perkin Elmer #CLS136334). Mice were maintained until day 70 or until humane endpoint as approved by UAB IACUC. For OVCAR8, 5×10⁵ mCherry^+^-, firefly luciferase^+^-OVCAR8 cells were injected subcutaneously in 50% Matrigel (Corning #356255). After 3 days, 1.5×10⁶ CAR^+^ T cells were injected IV. Tumor size was measured by calipers. Tumor volume was calculated as V = (1/2) * (Length * Width²). Mice were maintained until day 33 post-CAR-T cell infusion. Maximum tumor sizes were defined as 2000mm^3^, which was not exceeded by any mice in the study. Mice were not maintained past tumor-related symptoms, including significant ascites or decreased body condition. Mice were randomized to treatment groups once tumor burden was confirmed by ranking mice by tumor size and sequentially assigning one mouse per group. Each cage contained mice from at least two groups, limiting confounders. No mice were excluded once treatments were applied. Studies were conducted without blinding.

### *Ex vivo* analysis of CAR-T cells

Mice were perfused before tissue collection. Omental tumors were digested in IMDM supplemented with 5% bovine serum albumin, 2ng/mL Liberase TH (Roche #12352200), and 10U/mL DNase I (Thermo Scientific #EN0521). CAR-T cell numbers were quantified using Precision Count Beads (Biolegend #424902).

### Luminescence assay

NFAT-Luc transduced T cells were plated at 1×10^5^ cells per well, or Jurkat-NFAT-Luc cells were plated at 0.5×10^5^ cells per well in a 96-well round-bottom plate. Cells were stimulated overnight as indicated for each experiment, then lysed in a buffer containing 1% Triton X-100. Lysates were cleared by centrifugation before mixing with Luciferase-Assay Buffer containing 2mM adenosine triphosphate and 100µM Akalumine hydrochloride substrate (MedChemExpress #HY-112641A). The luminescence signal was measured with a Varioskan Lux Spectrophotometer.

### Cytokine Enzyme-Linked Immunosorbent Assay (ELISA)

2×10^5^ CAR-T cells enriched by flow sorting were incubated with the same number of Jurkat-HER2 cells in 200µL IMDM supplemented with 5% human AB serum for 48 hours in a 96-well round-bottom plate. Supernatants were collected after centrifugation at 2,000 x *g*. Analysis was performed through EVE Technologies (Alberta, Canada; #HD15).

### Western blotting

Western blots were performed as previously described^21^. Blots were developed with Pierce ECL Plus Western Blotting Substrate (Thermo Scientific #32134) and imaged for Alexa Fluor 488 with an iBright FL1500 imaging device (Invitrogen #A44241). Quantification was performed by densitometry in ImageJ. Antibodies used are listed in Supplementary Table 1.

### Statistics

Prism 10 (GraphPad, Boston, MA) was used for statistical analyses. Means are shown. Error bars represent the standard error of the mean (SEM). Specific tests used for each experiment are described in the figure legends. All statistical tests are two-tailed with a significant p<0.05. Statistical significance is depicted by: ns: p>0.05, *: p<0.05, **: p<0.01, ***: p<0.001, ****: p<0.0001.

### Ordinary differential equations (ODE) model of CAR-T cell therapy

Derivative functions were rationally designed based on the Lotka-Volterra model^22^ with inspiration from other CAR-T focused ODE models including CAR-T Math^23^. Equations were solved in R Studio using DeSolve^24^. Parameters were assumed or fit to the observed tumor curve by manual adjustment (see **Supplementary Table 2**). Per cell cytotoxicity rates were estimated using data from the cytotoxicity assay against OVCAR8 at E:T ratio of 2:1, with mathematical extrapolation for a per cell basis. Model fitting used the OVCAR8 tumor model with shJNK CAR-T cell treatment.

## Results

### JNK inhibition suppresses CD3-induced NFAT reporter activity

NFAT proteins are essential for T-cell activation and function^8^. To measure NFAT activity in CAR-T cells, we used Jurkat cells modified with an NFAT-Luciferase reporter (Jurkat-NFAT-Luc, **Supplemental Fig.1A**). Ionomycin stimulation induced >22-fold NFAT-Luc signal over unstimulated or PMA-stimulated controls (**Supplemental Fig.1B**), confirming NFAT responsiveness.

Based on reported interactions between JNK and NFAT^10–12^, we hypothesized that JNK inhibition impacts NFAT signaling in T cells. Pharmacologic JNK inhibitor SP600125^25^ suppressed NFAT reporter signal in Jurkat cells during αCD3/αCD28 antibody co-stimulation with a maximum decrease of 58% and a half maximal inhibitory concentration (IC50) of 2.75μM (**Supplemental Fig.1C**). Activated PBMC-derived T cells showed JNK and c-Jun phosphorylation upon αCD3/αCD28 antibody co-stimulation. SP600125 treatment inhibited JNK activity, resulting in decreased c-Jun phosphorylation and an over 80% decrease in NFAT reporter signal (IC50 of 7.8µM) (**Supplemental Fig.1D-G**). NFAT reporter signal suppression was independent of the degree of TCR activation or presence of αCD28 antibody (**Supplemental Fig.1H,I**), suggesting JNK responds to αCD3 antibody stimulation. The findings indicate that JNK inhibition reduces CD3-derived NFAT reporter activity in primary human T cells.

### JNK knockdown by shRNA suppresses CAR-induced NFAT signaling

We sought to manage NFAT activity via JNK suppression as a therapeutic strategy, but existing pharmacological JNK inhibitors are limited in their potency and specificity^26^. We designed an shRNA simultaneously targeting JNK1 and JNK2 (shJNK; sequence GATCATGAAAGAATGTCCTA) as an alternative. Lentivirally expressed shJNK knocked down JNK protein level in Jurkat-NFAT-Luc cells by 50%, and decreased NFAT reporter signal during αCD3/αCD28 antibody co-stimulation by 42%, comparable to 37% reduction by 10µM SP600125, (**Fig.1A-C**). These findings confirm that JNK activity enhances TCR-driven NFAT reporter activity, which can be reduced by specific knockdown or inhibition of JNK.

CAR signaling kinetics differ somewhat from those of natural TCRs^27^. We hypothesized that JNK knockdown suppresses NFAT reporter activity induced by CAR signaling. We constructed a HER2-targeting CAR using the single-chain form of Trastuzumab^28^, incorporating a CD28 spacer, transmembrane, and intracellular domain linked to CD3ζ. This CAR (HER2-28ζ) is lentivirally expressed with ZsGreen^29^ as a transduction marker (‘CAR-T cells’), or additionally co-expressed with the shJNK cassette (‘shJNK CAR-T cells’) (**Fig.1D**). In primary human CAR-T cells, shJNK reduced JNK protein levels by ∼50% (**Fig.1E,F**), and suppressed CAR-induced NFAT reporter activity by ∼40% after co-culture with Jurkat-HER2 cells (which were engineered to lack human leukocyte antigen (HLA) class I expression^30^, ensuring that CAR-T cell stimulation was not mediated through an HLA/TCR-dependent mechanism, **Fig.1G**). We concluded that JNK knockdown or inhibition suppresses CAR-induced NFAT reporter activity in primary human T cells.

### JNK knockdown does not hinder CAR-T cell production

We assessed how JNK knockdown influenced the characteristics of CAR-T cells post production. JNK knockdown did not affect the overall expansion of CAR-T cells or the CD4 to CD8 ratio (2.30 in control vs 2.57 in shJNK) (**Fig.1H-J**). Differentiation levels were also similar in the final product, mostly composed of CAR-T central memory (CAR-T_CM_: CD62L^+^/CD45RO^+^, control: 34%; shJNK: 36%) and CAR-T effector memory cells (CAR-T_EM_: CD62L^-^/CD45RO^+^, control: 36%; shJNK: 35%) (**Fig.1K,L**). Moreover, there were similar levels of three key transcription factor expressions for T-cell differentiation and function (TCF1, T-bet, and TOX)^31, 32^ (**Fig.1M,N**). Ultimately, JNK knockdown did not hinder CAR-T cell production or the differentiation profiles.

### JNK knockdown in CAR-T cells leads to improved cytotoxic activity

Since NFAT signaling supports the effector functions of T cells, including cytotoxicity in response to antigen engagement^7^, we sought to determine if dampening NFAT reporter activity via JNK suppression would impair CAR-T cell cytotoxicity. We compared the HER2-dependent killing against the human ovarian cancer cell lines OVCAR8 and SKOV3, which exhibit nearly 40-fold differential HER2 expression (mean fluorescent intensity (MFI) 274 vs 8103) (**Fig.2A,B**). Remarkably, shJNK CAR-T cells displayed a roughly two-fold increase in killing against both cell lines (**Fig.2C,D**). The evidence shows that JNK knockdown in CAR-T cells boosts their cytotoxic effectiveness against HER2^Hi^ and HER2^Lo^ ovarian cancer cells. This surprising discovery underscores the primary benefit of JNK knockdown in CAR-T cells: a direct increase in cytotoxicity.

**Fig. 2:**
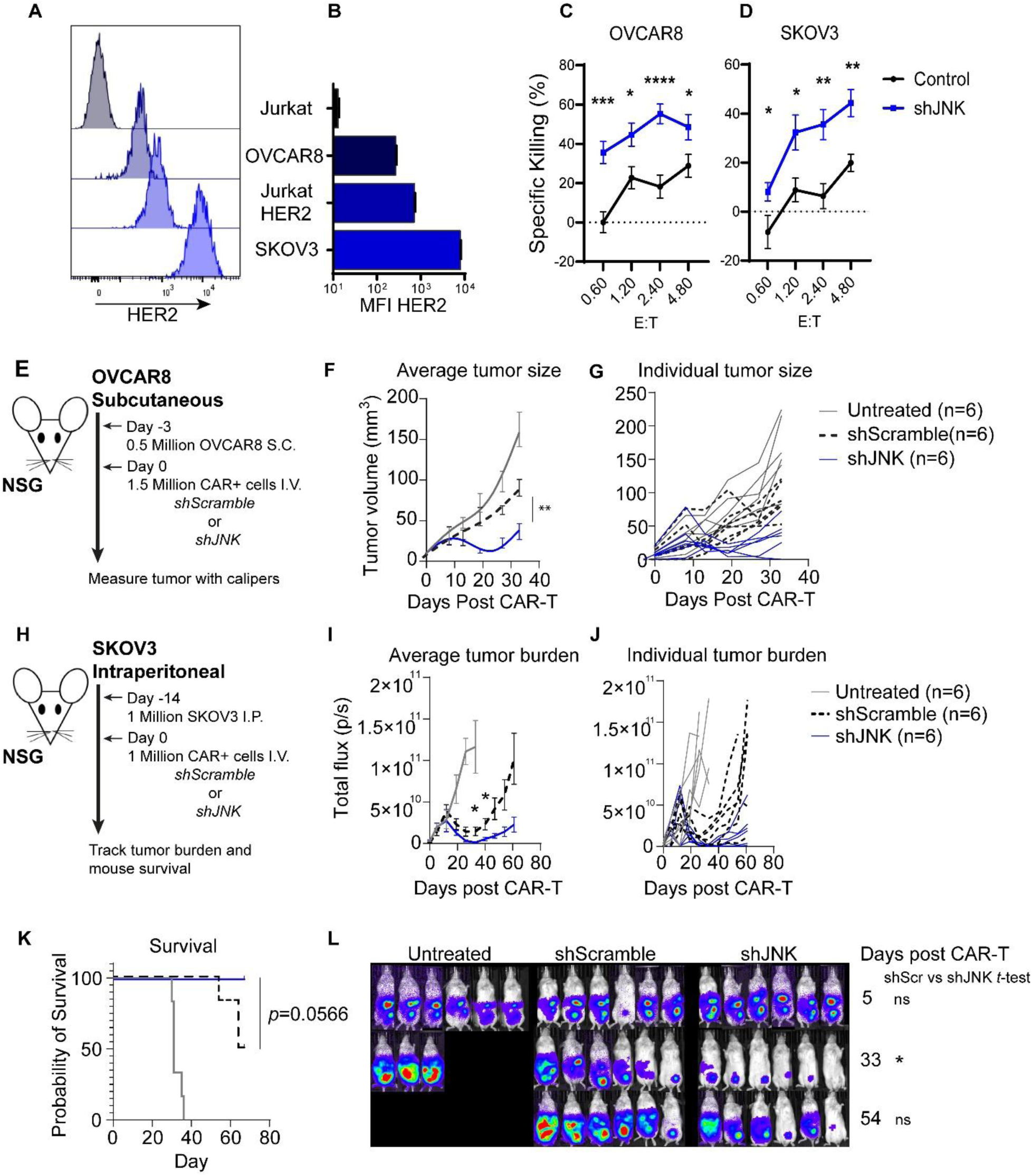
JNK knockdown CAR-T cell display enhanced cytotoxicity and *in vivo* efficacy. **A-B.** The indicated cell lines were analyzed for HER2 expression by flow cytometry; histograms (**A**) and MFI quantification (**B**). **C-D.** Cytotoxicity assays of Control vs. shJNK CAR-T cells against OVCAR8 and SKOV3 cells. N=3 donors, 4 technical replicates each. Aggregate data. **E-F.** Experimental Scheme (**E**). NSG mice were injected SC with 5×10^5^ OVCAR8 cells, followed by IV infusion of 1.5×10^6^ CAR^+^ T cells at day 3. Tumor burden was measured with calipers. 6 mice per group. One representative experiment of N=2 donors shown. Average (**F**) and individual (**G**) tumor volumes are shown. **H-L.** Experimental Scheme (**H**). NSG mice were injected IP with 1×10^6^ Firefly Luciferase^+^ SKOV3 cells, followed by IV infusion of 1×10^6^ CAR^+^ T cells at day 14. Tumor burden was assessed by bioluminescent imaging, with total flux (photons/second, p/s) used as a measure of tumor burden. 6 mice per group. One representative experiment of N=2 donors shown. Average (**I**) and individual (**J**) tumor burdens are measured by whole body BLI. Mouse survival (**K**) and representative BLI images (**L**) are shown. *p*-values are calculated using multiple unpaired *t*-tests with Welch’s correction. Survival curve statistics were calculated using Log-rank test.

### JNK knockdown CAR-T cells demonstrate superior anti-ovarian cancer efficacy *in vivo*

Since JNK knockdown enhanced CAR-T cell cytotoxicity (see **Fig.2C,D**) without impacting CAR-T cell production (see **Fig.1H-N**), we hypothesized JNK knockdown could enhance CAR-T cell performance *in vivo*. CAR-T cells with JNK knockdown or scramble shRNA sequence (shScramble) were evaluated against subcutaneous (SC) OVCAR8 tumors (**Fig.2E**), a High-Grade Serous Ovarian Cancer (HGSOC) type of tumor expressing a moderate level of HER2 and low levels of HLA class I molecules^33^. Starting on day 19, mice treated with shJNK CAR-T cells had significantly smaller tumors compared to those treated with shScramble CAR-T cells (**Fig.2F,G**). By day 33, shJNK CAR-T cells reduced tumor size by 77.5%, compared to 44.3% reduction by shScramble CAR-T cells. We further evaluated shJNK CAR-T cells in a model using firefly luciferase-labeled SKOV3 tumors engrafted IP (**Fig. 2H**), mimicking the abdominal metastasis seen in human ovarian cancer. At day 33, shJNK CAR-T-cell treated mice had 11.9-fold smaller tumor burden by BLI than the shScramble CAR-T treated mice (**Fig.2I-L**).

Notably, shJNK and shScramble CAR-T cells displayed similar response kinetics, with reduced tumor burden first seen at day 21 post-infusion and a peak response at day 33 against SKOV3 tumors. Because CAR-T cell exhaustion is a key barrier for durable anti-tumor responses^34^, and based on the similar response kinetics between shJNK and shScramble CAR-T cells, we hypothesized that JNK knockdown did not affect the development of CAR-T cell exhaustion during tumor challenge. We assessed the exhaustion status of CAR-T cells obtained from tumors of SKOV3-bearing mice through the expression of T-cell exhaustion markers (PD1, Tim3, Lag3, and TIGIT), and transcription factors (T-bet, TOX, TCF1, and Eomes) ^34, 35^. At day 21, CD8+-CAR-T cells from multiple donors exhibited both T-bet^+^/TOX^+^ and T-bet^-^/TOX^+^ phenotypes (**Supplemental Fig. 2A,B**). The presence of T-bet^+^ cells suggests retained functional activity^36^, consistent with reduced tumor burden at this time point, while T-bet^-^/TOX^+^ cells may indicate terminal exhaustion within the tumor environment. Notably, the analysis demonstrated that JNK knockdown did not result in any variations in the balance of these populations, the overall expression levels of transcription factors (**Supplemental Fig.2C**), or exhaustion marker expression (**Supplemental Fig.2D**). In addition, the surface expression of two stemness markers, CD62L and CC Chemokine Receptor 7 (CCR7)^17^, did not differ significantly between Control and shJNK CAR-T cells in three independent locations (**Supplemental Fig.2E,F**). These results demonstrate that JNK knockdown enhanced CAR-T cell anti-tumor efficacy, particularly by reducing tumor burden during the acute phase of treatment, but has no influence on T-cell stemness or the onset of T-cell exhaustion.

### Tumor-infiltrating JNK knockdown CAR-T cells display enhanced granzyme B expression

Based on enhanced cytotoxicity by JNK knockdown *in vitro*, and an enhanced ability to fight tumors *in vivo*, we sought to characterize the status of JNK knockdown CAR-T cells during tumor challenge. Specifically, we hypothesized that JNK knockdown CAR-T cells may show increased cytotoxic properties in the tumor environment. Twenty-eight days after infusion into SKOV3 tumor-bearing mice, shScramble and shJNK CAR-T cells showed similar biodistribution across the bone marrow, spleen, and omental locations, indicating that JNK knockdown does not affect *in vivo* expansion or homing (**Fig.3A**). Importantly, granzyme B (GZMB) expression was approximately doubled in CD8^+^-shJNK CAR-T cells within the omental tumor site (shJNK: 24.5%^+^, shScramble: 11.6%^+^; **Fig.3B**), with the increase only observed in the tumor environment. In a second donor experiment, we again observed similar biodistribution (**Fig.3C**). Although the increase in GZMB was not statistically significant due to small sample size (n=3), the trend toward higher GZMB in shJNK CAR-T cells persisted (average 48.2%^+^ in shJNK versus 22.5%^+^ in shScramble) and remained tumor-specific (**Fig.3D**). Other cytotoxic factors granzyme A (GZMA) and Fas Ligand (FASL) were not elevated by JNK knockdown (**Fig.3E,F**). Overall, these results support that JNK knockdown directly enhances tumor-localized GZMB expression in CD8^+^-CAR-T cells, without impacting the organ biodistribution of the CAR-T cells in the tumor-bearing mice.

**Fig. 3:**
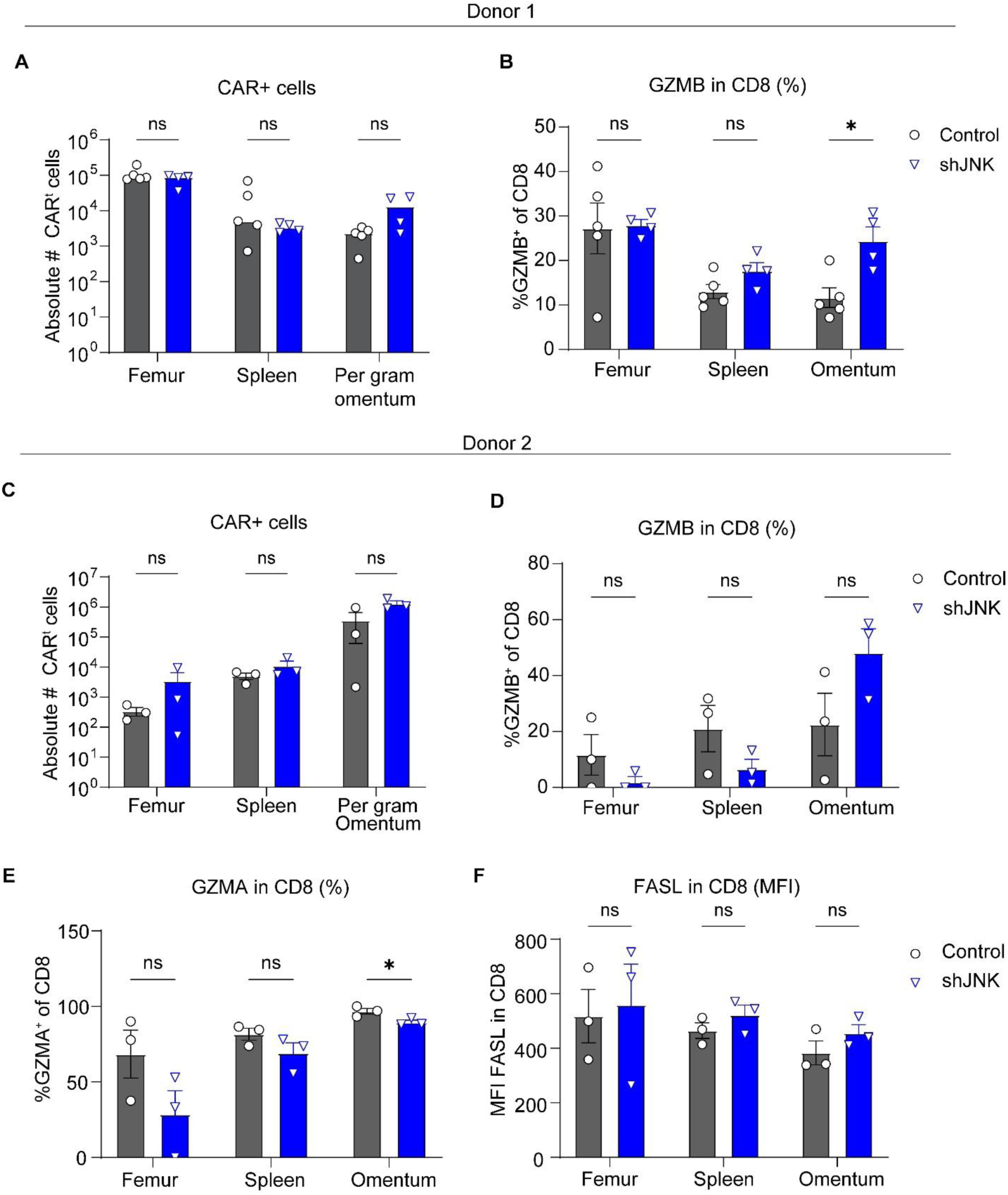
Tumor-infiltrating shJNK CAR-T cells display heightened GZMB expression. **A-B.** NSG mice were engrafted with 5 million Firefly Luciferase^+^-SKOV3 cells by IP injection, followed by I.V. infusion of 1 million CAR-T cells at day 14. Tissues were harvested 28 days after CAR-T cell infusion. Each data point represents one mouse. n = 5 mice control CAR-T, 4 mice shJNK CAR-T. Total number of CAR^+^-cells per tissue (**A**) and percentage of CD4^-^-CAR-T cells expressing GZMB (**B**) are shown. **C-F.** Ex vivo analysis of CAR-T cells as in (**A-B**), with a second donor. n = 3 mice control CAR-T, 3 mice shJNK CAR-T. Total number of CAR^+^ cells per tissue (**C**) and percentage of CD8^+^-CAR-T cells expressing GZMB (**D**) are shown. Additionally, percentage of CD8^+^-CAR-T cells expressing GZMA (**E**) and the MFI of FASL (**F**) on CD8^+^-CAR-T cells are shown. All *p*-values are calculated with multiple unpaired *t*-tests with Welch’s correction.

### JNK knockdown enhances CD8^+^-CAR-T cell killing and enhances degranulation activity

To test if the boost in GZMB levels is intrinsically linked to JNK knockdown rather than to secondary *in vivo* effects like differentiation or microenvironmental signals, we evaluated GZMB expression *in vitro* following overnight stimulation with recombinant HER2-Fc protein. JNK knockdown resulted in higher levels of GZMB, especially within the CD8^+^-subset, which showed a 1.6-fold increase in expression intensity (**Fig.4A**). Meanwhile, GZMA was also upregulated to a lesser extent than GZMB (1.4-fold increase in CD8^+^-CAR-T cells; **Fig.4B**) while FASL expression remained unchanged (**Fig.4C**). These findings reinforce the connection between JNK knockdown and elevated levels of cytotoxic factors, namely GZMB, with GZMA potentially playing a role in anti-tumor cytotoxicity *in vitro*. Importantly, we confirmed that JNK knockdown CAR-T cells display elevated GZMB levels after co-culture with OVCAR8 and SKOV3 cells (**Fig.4D-F**), with CD8^+^-shJNK CAR-T cells showing over 1.9-fold increase in GZMB intensity compared to controls after incubation with OVCAR8 cells. Since granzyme-mediated cytotoxicity depends on the release of secretory granules through T cell degranulation^37^, we measured CAR-T cell degranulation in response to HER2-Fc protein. JNK knockdown increased degranulation in both CD4^+^- and CD8^+^-CAR-T cells (**Fig.4G-I**), further supporting a general enhancement of the granzyme/degranulation pathway in both populations. However, when cytotoxicity was assessed separately using sorted CD4^+^- or CD8^+^-populations, only CD8^+^-CAR-T cells showed significantly increased killing activity after JNK knockdown (**Fig.4J-K**), highlighting the central role of CD8^+^-cells in this effect. Overall, the results demonstrate that JNK knockdown is directly linked to enhanced cytotoxicity of CD8^+^ CAR-T cells through the upregulation of key cytotoxic properties, including degranulation and granzyme expression.

**Fig. 4:**
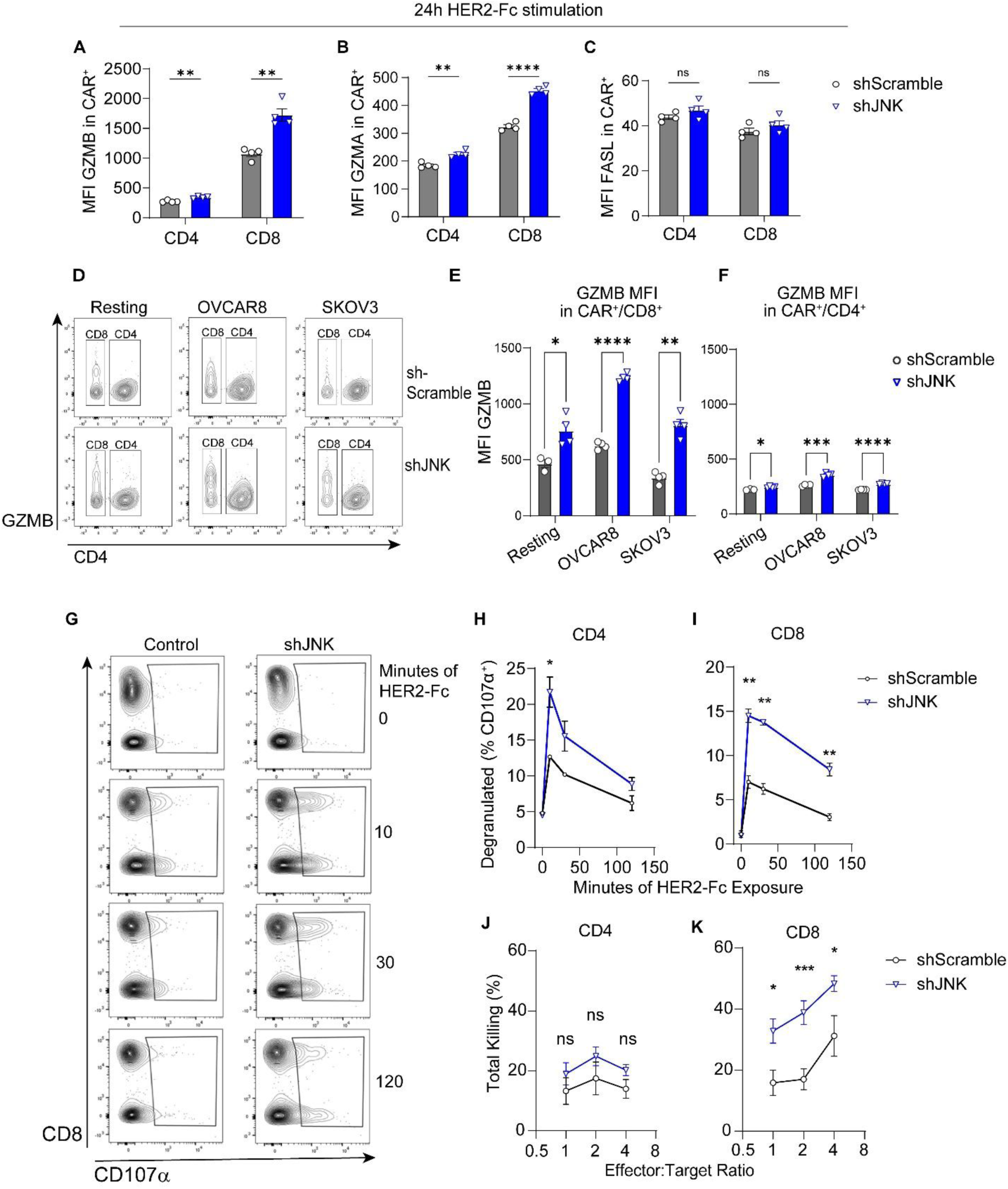
JNK knockdown enhances CD8^+^-CAR-T cell killing and enhances degranulation and granzyme activity. **A-C.** shScramble and shJNK CAR-T cells were incubated with 3µg/mL plate-bound HER2-Fc protein for 24 hours. Killing factors were assessed by flow cytometry. N= 1 representative experiment of 2 independent experiments with unique biological donors, 5 technical replicates each. Bar graphs show the MFI of GZMB (**A**) GZMA (**B**) and FASL (**C**) in CD8^-^ and CD8^+^ CAR-T cells. **D-F.** shScramble or shJNK CAR-T cells were co-cultured with 1×10^4^ OVCAR8 or SKOV3 cells for 24 hours before flow cytometry for GZMB. N = 1 experiment with 5 technical replicates shown. Multiple unpaired *t-*tests with Welch’s correction. Representative flow panels (**D**) depict the populations with MFI quantifications in (**E-F**). **G-I.** shScramble or shJNK CAR-T cells were incubated with 3µg/mL plate-bound HER2-Fc protein for the indicated time, then monensin and αCD107α and αCD8 antibodies were added. Degranulation was assessed by CD107α^+^ in flow cytometry; representative plots (**G**). Degranulation was quantified in CD8^-^-(**H**) and CD8^+^-(**I**) CAR-T cells. N = 2 experiments with unique biological donors. One experiment shown. 3 technical replicates per group. **J-K.** Sorted CD4^+^-CAR^+^ (**J**) or CD8^+^-CAR^+^ (**K**) T cells were used for cytotoxicity assay against OVCAR8 cells. Total killing shown. N = 2 independent experiments with unique biological donors, 4 technical replicates each. Aggregated data. All *p*-values calculated using multiple unpaired *t-*tests with Welch’s correction.

### JNK knockdown promotes GZMB upregulation across phenotypes and is linked to NFATc1 activity

As effector CD8^+^-T cells are thought to possess the highest level of cytotoxic activity^38^, we considered that the enhanced cytotoxicity observed in JNK knockdown CAR-T cells might arise from selective activation of specific memory or effector populations, rather than a uniform increase in cytotoxic potential. To address this, we assessed the contribution of distinct CAR-T cell phenotypes to the upregulated GZMB levels following JNK knockdown. Under αCD3/αCD28 antibody co-stimulation, JNK knockdown enhanced intensity of GZMB expression-nearly threefold-in CD8^+^ CAR-T cells compared to controls (MFI 1750 vs MFI 658, **Fig.5A**), with similar increases across CAR-T stem-cell memory (CAR-T_SCM_)/ CAR-T naïve (CAR-T_N_), CAR-T_CM_, CAR-T_EM_, and CAR-T effector (CAR-T_EFF_) populations (**Fig.5B,C**). These findings indicate that the upregulation of GZMB following JNK knockdown is independent of the CAR-T cell differentiation state.

**Fig. 5:**
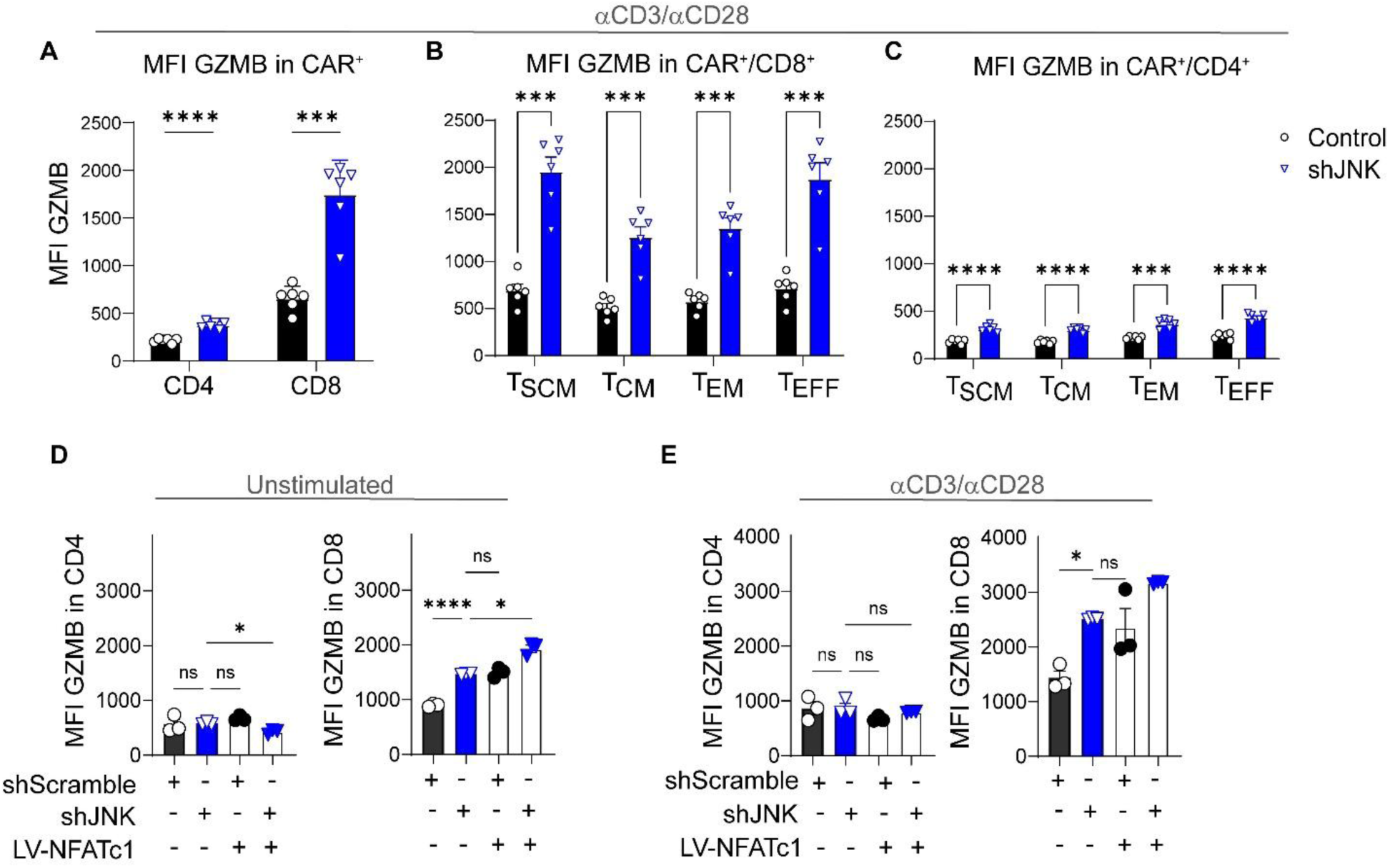
JNK knockdown elevates GZMB across phenotypes and is linked to NFATc1 activity. **A-C.** shScramble or shJNK CAR-T cells were stimulated with 2.5% w/v Immunocult CD3/CD28 antibody complexes for 24 hours before analysis of GZMB expression by flow cytometry. Distinct populations are segregated by CD62L and CD45RO expression. N = 2 donors. One representative experiment with 6 technical replicates shown. Multiple unpaired *t-*tests with Welch’s correction. The overall MFI of GZMB (**A**), as well as the GZMB MFI in subpopulations of CD8 (**B**) and CD4 (**C**) CAR-T cells is shown. **D-E.** Primary human CAR-T cells were transduced with lentiviral vectors encoding NFATc1. Representative flow plots show GZMB expression after 2.5% w/v Immunocult CD3/CD28 antibody stimulation. N = 2 independent experiments with one donor. One representative experiment shown with 3 technical replicates without (**D**) or with (**E**) 2.5% w/v Immunocult CD3/CD28 antibody stimulation. Brown-Forsyth and Welch ANOVA.

The ultimate question is the molecular mechanism connecting JNK knockdown and enhanced cytotoxic activity. Since JNK activation inhibits NFATc1^10, 12^, which is reported to promote GZMB expression in murine T cells^7^, we hypothesized that JNK knockdown enhances GZMB expression through NFATc1. We overexpressed NFATc1 in CAR-T cells, hypothesizing that changing the NFATc1/JNK balance would be sufficient to enhance GZMB expression. Overexpression of NFATc1 in control CD8^+^-CAR-T cells elevated the GZMB levels to a similar degree as JNK knockdown (**Fig.5D,E**), suggesting that the balance between NFATc1 and JNK levels is essential for the level of GZMB expression. Overall, the results imply that the improved tumor control *in vivo* following JNK knockdown is linked to increased cytotoxicity, driven by elevated GZMB levels in the tumor through an NFATc1-dependent mechanism.

### JNK suppression alters multiple outcomes of T cell stimulation

JNK knockdown increased the level of GZMB expression, which we suggest is mediated through enhanced activity of NFATc1. However, JNK knockdown also suppressed NFAT reporter activity, representing a paradoxical result. Based on a literature report that JNK enhances NFATc2 transcriptional activity^11^, we hypothesized that JNK knockdown may selectively enhance cytotoxic responses through NFATc1 activation while suppressing other activation pathways through NFATc2 suppression. We used surface expression of CD25 and CD69 as T-cell activation markers, as their expression is impacted by NFAT activity^39^. JNK inhibition during stimulation of PBMC-derived T cells reduced CD25 and CD69 expression by ∼45% (**Supplemental Fig.3A-D**). The suppression was broadly consistent across differentiation stages (**Supplemental Fig.3E-G**) and between CD4^+^- and CD8^+^-T cells (**Supplemental Fig.3H-I**). When shJNK CAR-T cells were stimulated through interaction with Jurkat-HER2 cells, consistent reductions in CD25 expression were observed (**Fig.6A-C**). The results indicate that JNK plays a role in the stimulation triggered by both TCR and CAR, affecting naïve and memory cells isolated from the blood as well as previously activated CAR-T cells. Using a multiplex ELISA assay, we further assessed CAR-T cell cytokine production as a major outcome of CAR T cell activation^40^ (**Fig.6D**). JNK knockdown reduced some cytokines (IL-10, IL-4, IL-5, Il-2) by roughly half and produced a trend toward reduction of Tumor necrosis factor alpha (TNFα). IL-6 and Interferon gamma (IFNγ) - which is critical for maintaining cytotoxic anti-tumor responses^41^ – were unaffected by JNK knockdown. Effects on Granulocyte-Macrophage Colony Stimulating Factor (GM-CSF), IL-13, and IL-8 varied between donors. Overall, while the cytotoxic response was enhanced, JNK knockdown in CAR-T cells also suppressed multiple facets of the antigen-stimulation response, consistent with reductions in NFAT reporter activity.

**Fig. 6:**
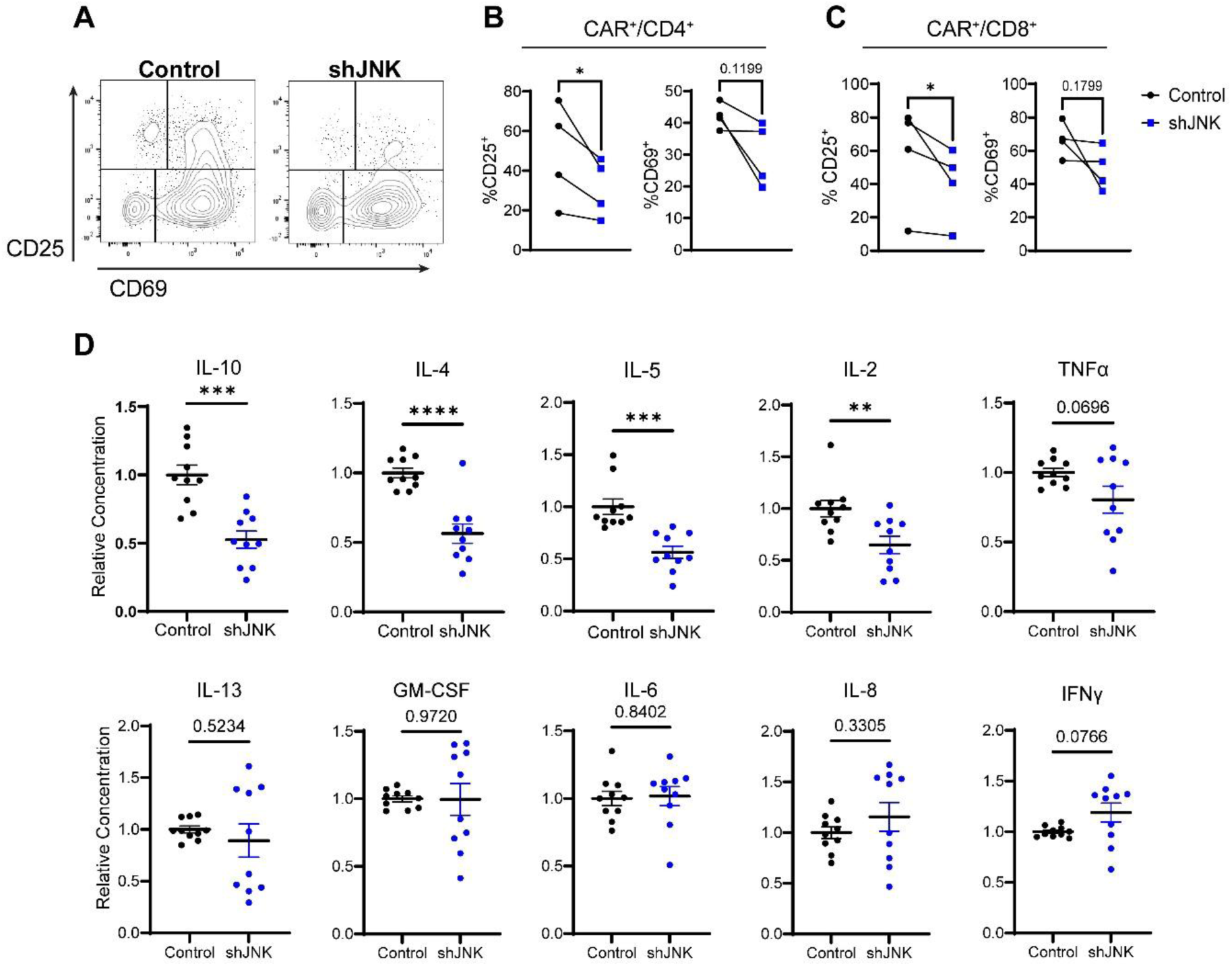
JNK knockdown suppresses antigen mediated CAR-T cell stimulation. **A-C.** Control and shJNK CAR-T cells were co-cultured with Jurkat-HER2 cells for 36 hours, then assessed for CD69 and CD25 expression by flow cytometry. N=4 donors. 2-6 technical replicates each. Different donors used different flow panels, with one representative flow plot shown in (**A**). Quantification of CD69 and CD25 expression on CD4^+^-(**B**) or CD8^+^-T cells (**C**) plotted as the average of each donor. Ratio Paired *t-*tests. **D.** Control and shJNK CAR-T cells were co-cultured with Jurkat-HER2 cells for 48 hours. Supernatant was harvested and used for multiplex ELISA analysis of cytokine production. Scatter plots show relative cytokine concentration produced by shJNK CAR-T cells compared to control CAR-T cells. Aggregated data of N=2 donors, 5 technical replicates each. Unpaired *t-*test with Welch’s Correction.

### Mathematical modeling suggests that the direct enhancement of cytotoxicity explains enhanced tumor control

Though JNK knockdown induced wide-ranging impacts on CAR-T cell stimulation, we hypothesized that enhanced cytotoxicity is sufficient to explain superior anti-tumor efficacy by shJNK CAR-T cells. We developed an ordinary differential equations (ODE) model of CAR-T cell therapy based on the Lotka-Volterra predator/prey interaction model^22^ (**Fig.7A**). Our model is unique in that it accounts for separate compartments of blood localized versus tumor localized CAR-T cells, CAR-T persistence in the blood, the rate of tumor infiltration, and an exhaustion like decay in cytotoxic function (**Fig.7B**). Parameter estimation is summarized in (**Supplementary table 2**). Briefly, tumor cell numbers were inferred from observed tumor size, CAR-T cell cytotoxicity was determined empirically by *in vitro* assay, and other parameters were fit using the observed tumor curves in untreated and shJNK treated mice (see: **Fig.2E-G**). Validating the principal of the model, we were able to simulate a dose dependent response to CAR-T cell infusion (**Fig.7C**). Testing our hypothesis, we kept all parameters equal, but adjusted the cytotoxicity to match the observed cytotoxicity *in vitro* (shScramble: 1.3–1.87, shJNK: 2.77-2.96 tumor cells killed/CAR-T cell/day) and ran 4 simulations. This difference in cytotoxicity produced significantly different tumor sizes in the model at day 33 (**Fig.7D**). Importantly, the simulated results closely aligned with the observed tumor growth, with no significant difference seen between simulated and observed tumor sizes on day 33 for any treatment group. Overall, our model demonstrated that the observed difference in cytotoxicity *in vitro* between control and shJNK CAR-T cells is sufficient to explain the different anti-tumor efficacy *in vivo*. This suggests that enhanced functionality of JNK knockdown CAR-T cells is primarily mediated through enhanced cytotoxicity.

**Figure 7:**
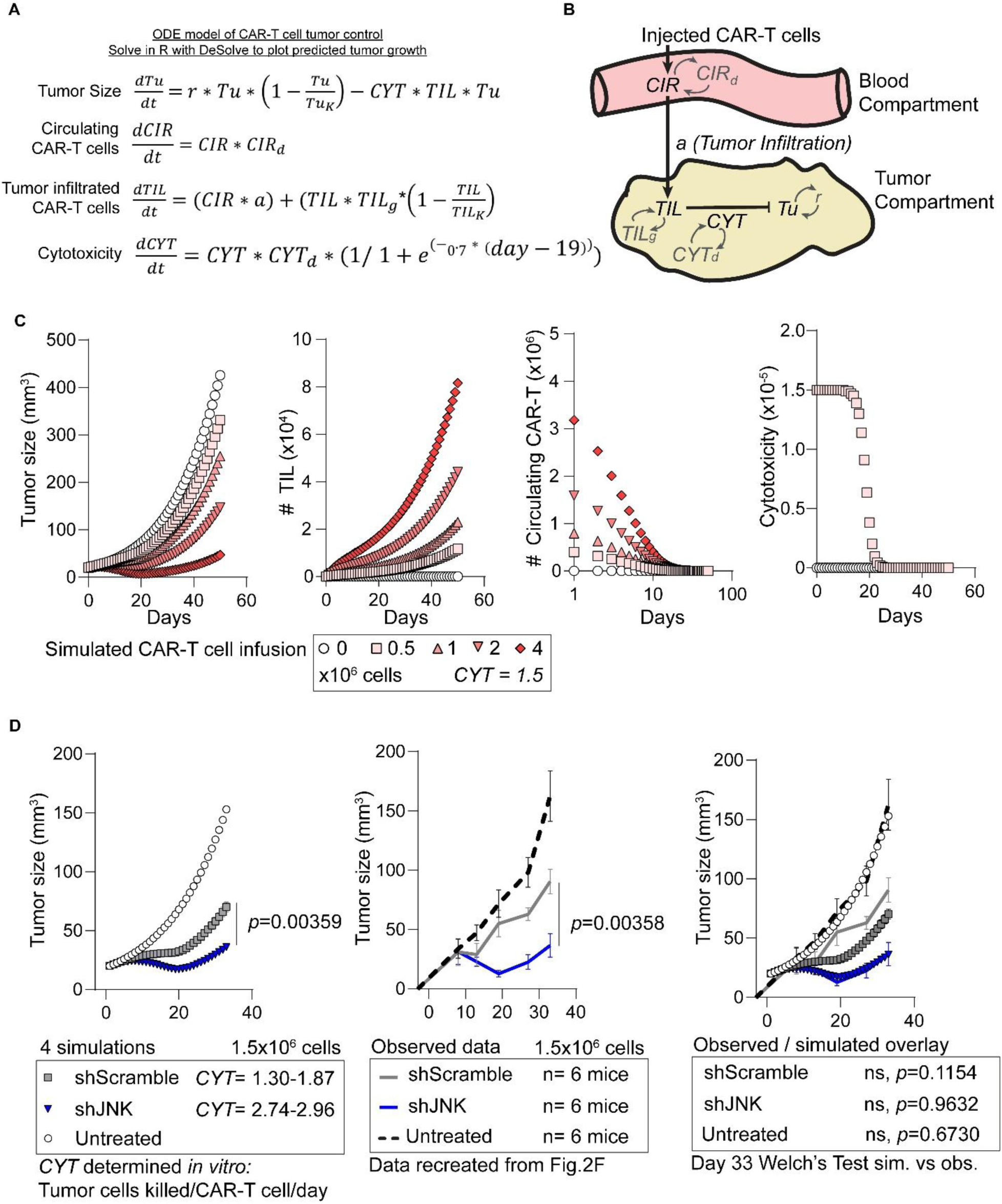
Mathematical modeling supports enhanced cytotoxicity as a mechanism for enhanced efficacy by JNK knockdown. **A.** Ordinary differential equations model of CAR-T cell therapy based on lotka-volterra interaction. Differential equations are shown with a label describing their function. Equations are solved using DeSolve in R to generate virtual tumor size plots. *CYT* = cytotoxicity, *Tu* = tumor size, *r* = tumor growth rate, *TIL* = number of tumor infiltrated CAR-T cells, *CIR* = number of circulating CAR-T cells, *a* = rate of tumor infiltration for circulating CAR-T cells, *X_d_* = decay of ‘x’, *X_g_* = growth of ‘x’, *X_k_* = carrying capacity of ‘x’. **B.** Diagram depicting interactions between model parameters. **C.** Model demonstration. After fitting the model using observed tumor growth for untreated and shJNK treated mice, tumor growth was simulated using the indicated doses of CAR-T cells. **D.** Cytotoxicity rates observed *in vitro* for shScramble and shJNK CAR-T cells were used to generate four simulations of tumor growth. These simulated tumor growth curves are overlaid with observed growth curves from (**Fig.2F**). Unpaired *t*-tests with Welch’s correction.

## Discussion

Our study found that JNK knockdown in CAR-T cells reduced overall CAR stimulation while increasing cytotoxicity and granzyme expression. Unlike most previously reported methods for enhancing CAR-T cell efficacy^1–5^, JNK knockdown did not affect CAR-T cell production, biodistribution, stemness, or exhaustion, highlighting the safe application of its therapeutic potential in CAR-T cell therapies.

JNK knockdown in CAR-T cells produced a robust, approximately two-fold increase in cytotoxic activity *in vitro*, which corresponded closely with a similar elevation in GZMB expression and degranulation. Enhanced cytotoxicity was constrained to CD8^+^-CAR-T cells, which showed the highest level of GZMB expression compared to CD4^+^-cells. Our findings support a model in which JNK knockdown primarily enhances GZMB gene expression in cytotoxic cells, yet fails to overcome potential barriers (for example, epigenetic barriers or transcriptional repressors) present in non-cytotoxic populations. Additionally, our analyses showed that exhaustion mechanisms remained intact after JNK knockdown, indicating the CAR-T cells retain the intrinsic regulatory controls that limit aberrant activation and toxicity^34^.

Our results align with a study in OT-I T cells showing that JNK2 deficiency elevates GZMB and cytotoxicity^42^. However, a parallel study found that JNK1 deficiency impairs initial OT-I cell activation^43^. Crucially, we used pre-activated human CAR-T cells, meaning JNK knockdown was introduced post-priming. In our case, JNK knockdown did not alter the differentiation properties of the CAR-T cells after production, and the stemness and exhaustion properties were not changed by JNK knockdown during *in vivo* tumor challenge. Thus, JNK appears to restrain CD8^+^-CAR-T cell cytotoxicity during the effector phase without influencing cell fate decisions.

JNK regulates multiple essential pathways, including activator protein 1 (AP-1) through interactions with c-Jun and other molecules^44^. However, our data indicate that JNK knockdown enhances CAR-T cell cytotoxicity through the NFAT pathway rather than AP-1 signaling. First, JNK knockdown in Jurkat cells did not alter p-c-Jun protein levels (data not shown), which is consistent with unaltered p-c-Jun levels under JNK2 deficiency^42^, suggesting AP-1 is not the primary mediator of altered GZMB levels or NFAT reporter levels. Second, a prior work has shown that c-Jun overexpression can mitigate CAR-T cell exhaustion^45^. Still, our results indicated that JNK knockdown selectively boosts cytotoxicity without affecting the exhaustion or differentiation status. JNK suppresses NFATc1 activity directly^10, 12^, and NFATc1 drives GZMB expression and granule polarization in murine CD8^+^-T cells^7^. NFATc1 overexpression was sufficient to recreate elevated GZMB levels similar to JNK knockdown. Therefore, we propose that JNK knockdown enhances CAR-T cytotoxicity by relieving inhibition on NFATc1, which then directly promotes GZMB transcription and degranulation. However, direct confirmation of NFATc1 and AP-1 activity after JNK knockdown are needed to confirm this mechanism.

This study is limited in that it did not determine direct molecular contacts between JNK and NFAT isoforms, nor rule out broader effects of JNK on T-cell activation (such as alternative splicing^46^ or autophagy^47^) that could indirectly enhance cytotoxicity. Thus, while a direct NFATc1-dependent mechanism is our leading model, off-target influences, changes in feedback loops, or involvement of other transcriptional regulators cannot be excluded. Further biochemical and genomic studies would be needed to evaluate these detailed mechanisms.

Most strategies to enhance CAR-T cell efficacy target differentiation, exhaustion, or biodistribution^1–5^, indirectly increasing overall cytotoxicity by improving CAR-T cell persistence and tumor infiltration. Meanwhile, direct methods to boost killing are limited; for example, one study overexpressed granzymes directly, which improved cytotoxicity but increased CAR-T cell apoptosis and limited *in vivo* persistence^48^. In our study, JNK knockdown enhanced CAR-T cell cytotoxicity without impacting *in vivo* persistence, which substantially benefits *in vivo* tumor killing. However, durable tumor control was not achieved in either model tested. Our *ex vivo* analysis of CAR-T cells suggested that they are still susceptible to exhaustion after JNK knockdown, likely explaining a loss of anti-tumor function after day 33. This emphasizes the need for strategies that also support CAR-T cell longevity. Combining JNK knockdown with such strategies may yield synergistic benefits for solid tumor therapy, though further investigation is needed to determine compatibility.

Beyond direct tumor cell killing, the improved cytotoxic function of JNK knockdown CAR-T cells may also influence the tumor microenvironment; increased cytotoxicity can promote tumor antigen release, and inflammation could be further amplified by the observed reduction in Th2 cytokines (IL-4, IL-5, IL-10) along with the preservation of IFNγ production^49^. These changes could potentially enhance antigen presentation and host immune responses by promoting the so-called cancer immunity cycle^50^. However, our current study was limited to xenograft systems. One future direction of the project is to evaluate these effects in syngeneic murine and humanized models of ovarian cancer.

## Conclusions

This study indicates that JNK knockdown boosts CAR-T cell effectiveness by increasing cytotoxicity. This is highly significant since it can considerably enhance the anti-cancer performance of CAR-T cells by just incorporating the shJNK expression cassette into any CAR vectors in use. It might also be compatible with existing methods that strengthen CAR-T cells’ longevity and stem-like characteristics, leading to enduring tumor eradication. Such improved functionality could mitigate the existing issues with CAR-T effectiveness in various solid tumors.

## Declarations

## Ethics statement

The University of Alabama at Birmingham Institutional Animal Care and Use Committee (IACUC) reviewed and approved the animal study.

## Consent for publication

Not applicable.

## Data availability statement

The original contributions presented in the study are included in the article. The corresponding authors can be contacted for further inquiries of source data.

## Author contributions

Conceptualization: CJK and MK, methodology: CJK and MK, formal analysis: CJK and MK, investigation: CJK, CEJ, MTB, FS, AM, and YNK. Writing: CJK and MK.

## Acknowledgments

Animal imaging was performed in the Small Animal Imaging Shared Facility at UAB supported by the O’Neal Comprehensive Cancer Center Grant P30CA013148. Flow cytometry was performed in the Flow Cytometry and Single Cell Core Facility at UAB supported by the Center for AIDS Research grant AI027767 and P30CA013148.

## Abbreviations

CAR: Chimeric Antigen Receptor
NFAT: Nuclear Factor of Activated T cells
JNK: c-Jun N-terminal Kinases
shRNA: short hairpin ribonucleic acid
HER2: Human Epidermal Growth Factor Receptor 2
TCR: T-cell receptor
IMDM: Isocove’s modification of Dulbecco’s medium
α/α: antibiotic/antimycotic solution
NFAT-Luc: Lentiviral vector responsive to NFAT reporter activity with luciferase expression
ARRE-2: Antigen Receptor Response Element 2
PBMC: Peripheral Blood Mononuclear Cells
LRS: Leukocyte retention system
PBS: Phosphate buffered saline
RBCs: Red blood cells
E:T: Effector:Target Ratio
PFA: Paraformaldehyde
IACUC: Institutional animal care and use committee
NSG: NOD.Cg-Prkdcscid Il2rgtm1Wjl/SzJ
IP: Intraperitoneal
IV: Intravenous
BLI: Bioluminescent intensity
PMA: Phorbol 12-myristate 13-acetate
ELISA: Enzyme linked immunosorbent assay
SEM: Standard Error of the Mean
ODE: Ordinary Differential Equations
IC50: Half Maximal Inhibitory Concentration
shJNK: short hairpin RNA targeting JNK1/2
HER2-28ζ: CAR construct consisting of HER2 binding domain, CD28 spacer, transmembrane, and linker, and CD3ζ intracellular domain
HLA: Human leukocyte antigen
CAR-T_CM_: CAR-T central memory cells
CAR-T_EM_: CAR-T effector memory cells
TCF-1: T-cell Factor 1
T-bet: T-box Expressed in T cells
TOX: Thymocyte Selection Associated High Mobility Group Box
MFI: Mean fluorescent intensity
shScramble: short hairpin RNA with the same bases as shJNK, but in a random order
SC: Subcutaneous
HGSOC: High Grade Serous Ovarian Cancer
GZMB: Granzyme B
GZMA: Granzyme A
FASL: Fas ligand
PD1: Programmed cell death protein 1
TIM3: T-cell immunoglobulin and mucin domain-containing protein 3
LAG3: Lymphocyte Activation Gene 3
TIGIT: T cell immunoreceptor with immunoglobulin and ITIM domain
CCR7: CC chemokine receptor 7
CAR-T_SCM_: CAR-T stem cell memory cells
CAR-T_N_: CAR-T naïve cells
CAR-T_EFF_: CAR-T effector cells
GM-CSF: Granulocyte-Macrophage Colony Stimulating Factor
IFNγ: Interferon Gamma
TNFα: Tumor Necrosis Factor alpha
AP-1: Activator protein 1

