## Supplemental information for "JNK knockdown enhances CAR-T cell cytotoxicity"

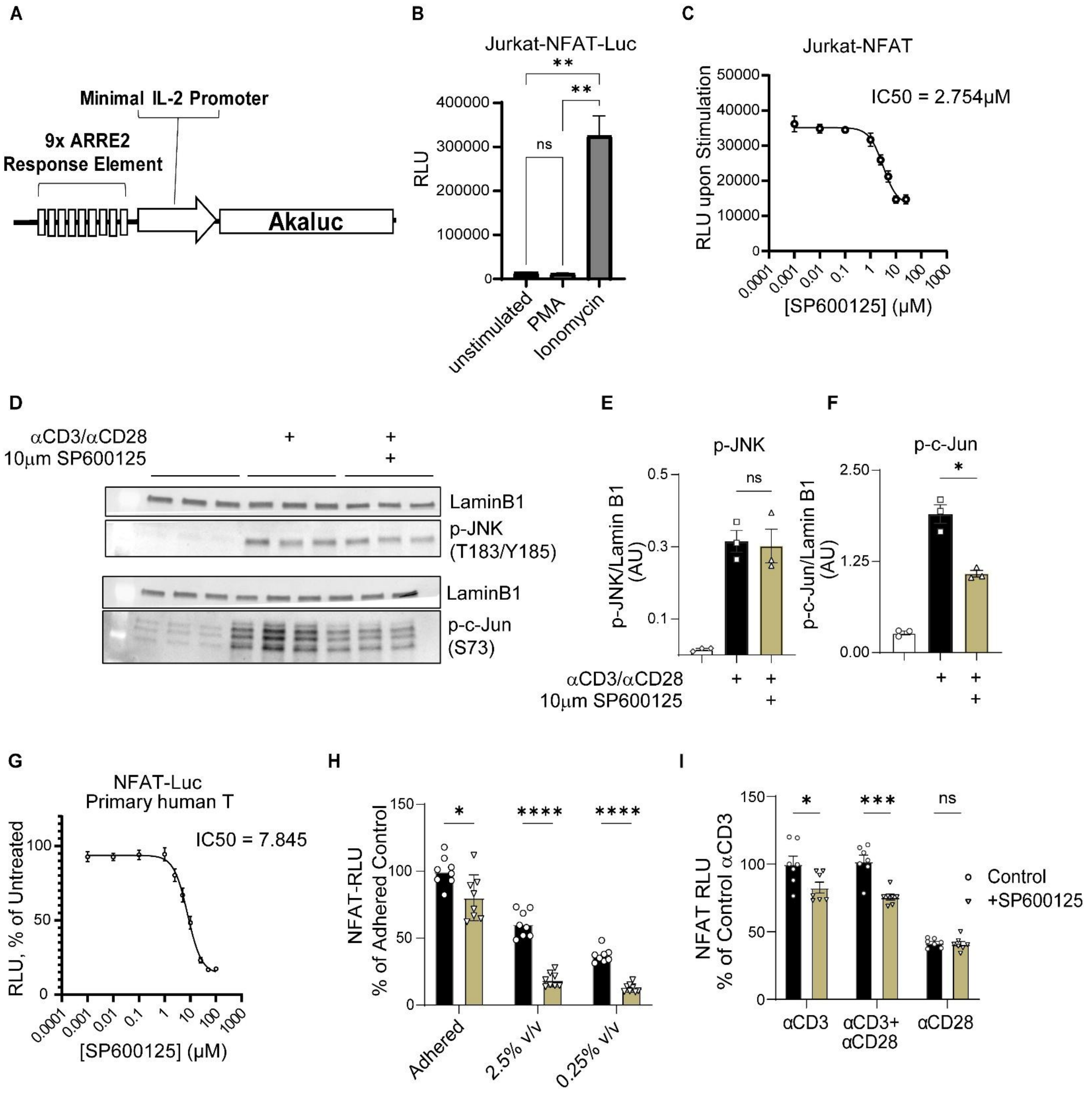


**Supplemental Fig.1: Impact of SP600125 on NFAT reporter activity.**

**A.** Schematic showing NFAT-Luc Reporter.

**B.** Luminescence assay of Jurkat-NFAT-Luc cells. One representative experiment with 4 technical replicates per condition.

**C.** Luminescence assay of Jurkat-NFAT-Luc cells with αCD3/αCD28 antibody co-stimulation and indicated SP600125 concentration. One representative experiment with 4 technical replicates per concentration.

**D-F.** Western blot and quantification of phospho-JNK and phospho-c-Jun in activated primary human T cells with 2-hour αCD3/αCD28 stimulation +/- SP600125. N=2 donors, three technical replicates. One representative experiment shown.

**G.** Luminescence assay of NFAT-Luc modified primary human T cells stimulated with αCD3/αCD28 antibody co-stimulation +/- SP600125 at the indicated concentration. Data is normalized to NFAT RLU in untreated (0μM SP600125) wells. N= 3 donors, 4 technical replicates. Aggregated data.

**H.** Luminescence assay of NFAT-Luc modified primary human T cells with αCD3/αCD28 antibody co-stimulation at 2.5% v/v adhered onto the plate, 2.5% v/v soluble, or 0.25% v/v soluble +/- 10μM SP600125. N=2 donors, 4 technical replicates. Aggregated data.

**I.** Luminescence assay of NFAT-Luc modified primary human T cells with αCD3, αCD28, or αCD3 + αCD28 antibodies +/- 10μM SP600125. N=1 representative experiment with 6 technical replicates per condition.

Statistical tests: Brown-Forsythe and Welch ANOVA with Dunnett's T3 (**B**), unpaired *t*-tests with Welch's correction (**E,F,H,I**).

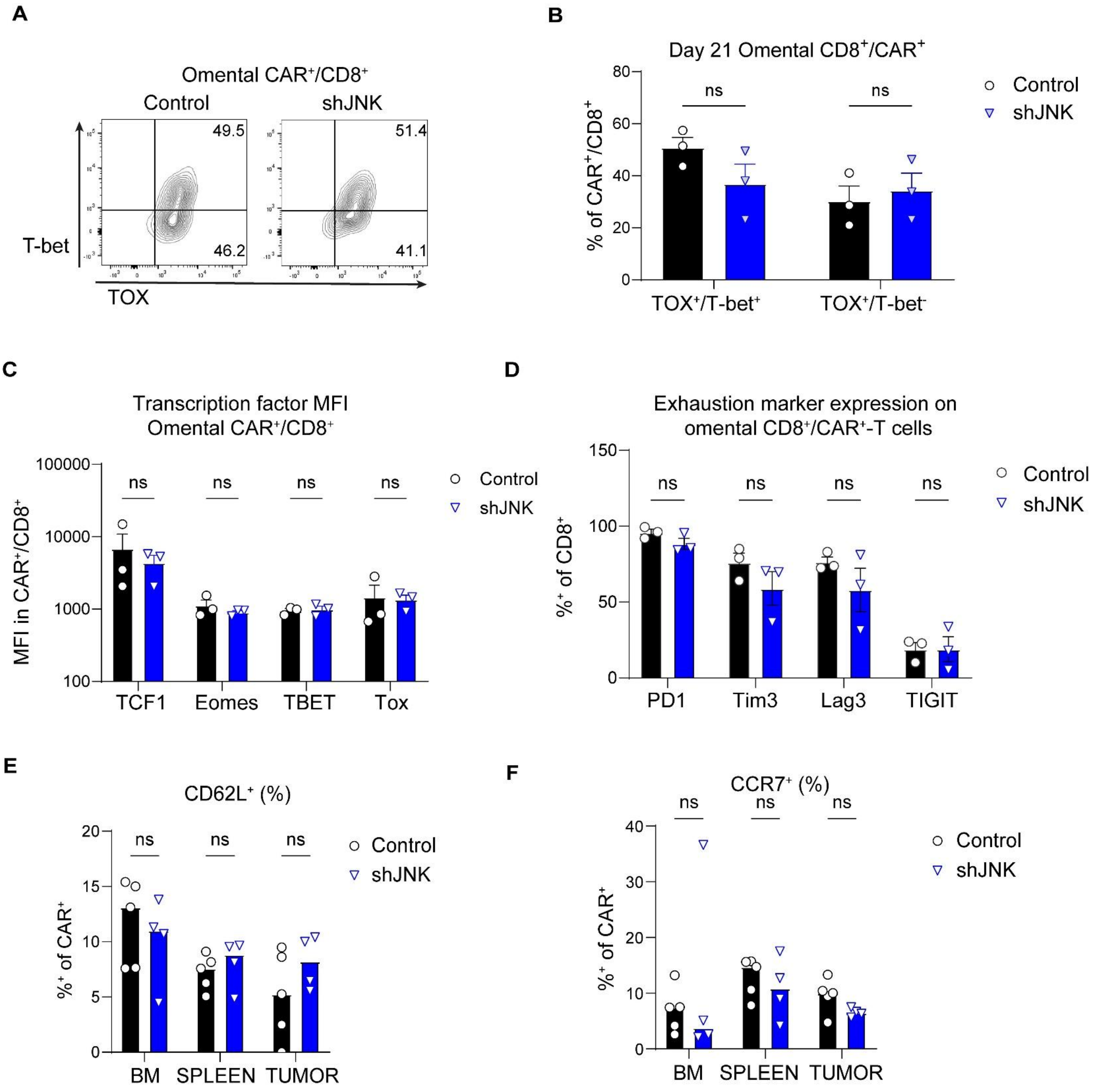


**Supplemental Fig.2: JNK knockdown CAR-T cells do not show altered exhaustion or differentiation status in vivo.**

**A-D.** NSG mice were engrafted with 5 million Firefly Luciferase⁺-SKOV3 cells via I.P. injection, followed by IV infusion of 1 million CAR-T cells at day 14. Mice were sacrificed at day 21 post-infusion and CAR-

T cells were analyzed by flow cytometry. Representative flow plots (**A**) show T-bet and TOX co-expression, which is quantified in (**B**). Total transcription factor MFIs are shown in (**C**). Exhaustion markers were stained separately, bar plot (**D**) shows the percentage of $CD8^+$-cells expressing each marker. **A-C**, N=2 independent experiments with unique biological donors. One representative experiment shown. **D**, N**=**2 independent experiments with unique biological donors, with the second experiment analyzing cells at day 14 post infusion. One representative experiment shown.

**E-F.** CAR-T cells day 28 post infusion. N = 2 independent experiments with unique biological donors. Representative experiment shown. Bar plots showing CD62L (**E**) and CCR7 (**F**) expression on $CAR^+$ cells in indicated tissues. BM = Bone marrow.

Each data point represents one mouse. *p*- values were calculated by multiple unpaired *t*-tests with Welch's correction.

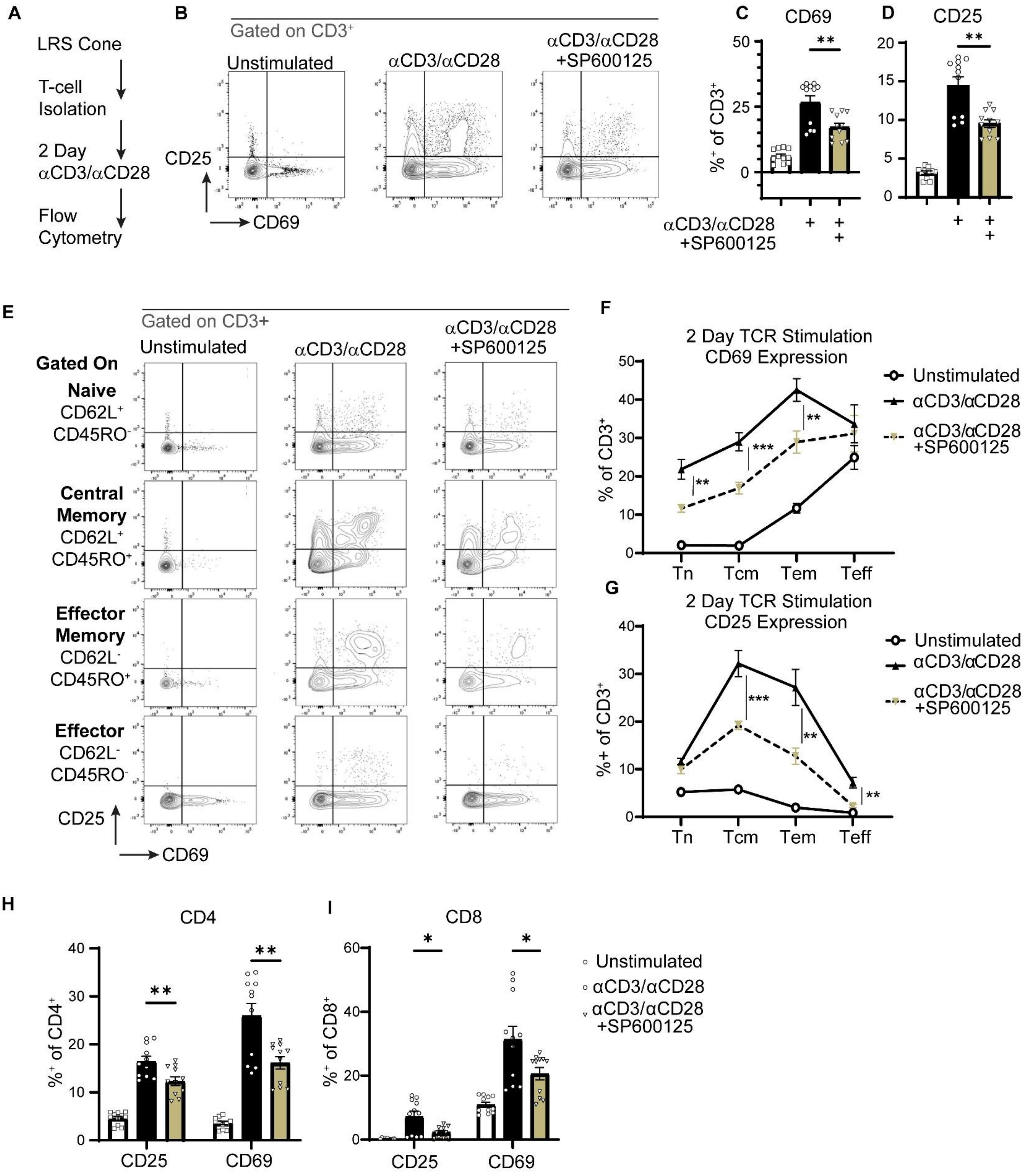


**Supplemental Figure 3: JNK inhibition suppresses TCR stimulation across T cell phenotypes.**

*Kuhlmann et. al. JNK knockdown enhances CAR-T cell cytotoxicity. Supplement.*

**A.** Experimental scheme for (**B-I**). Primary human T cells were isolated from PBMCs and incubated with 2.5% w/v Immunocult CD3/CD28 antibody stimulation for 2 days before analysis of CD69 and CD25 expression by flow cytometry. N = 3 donors, 3 technical replicates. Aggregated data.

**B.** Flow plots showing CD25 and CD69 expression on Live/CD3$^+$ cells.

**C.** Quantification of CD69 expression as in (**B**). Unpaired *t*-test with Welch's Correction.

**D.** Quantification of CD25 expression as in (**B**). Unpaired *t*-test with Welch's Correction.

**E.** Cells from (**A**) and (**B**) were segregated into distinct phenotypes by CD62L and CD45RO expression, flow plots for CD69 and CD25 in each indicated population are shown.

**F.** Quantification of CD69 expression in each subset as in **(E)**. Multiple Unpaired *t*-test with Welch's Correction.

**G.** Quantification of CD25 expression in each subset as in **(E)**. Multiple Unpaired *t*-test with Welch's Correction.

**H.** Quantification of CD69 and CD25 expression on CD4$^+$-T cells as in **(B)**. Multiple unpaired *t*-tests with Welch's Correction.

**I.** Quantification of CD69 and CD25 expression on CD8$^+$-T cells as in **(B)**. Multiple unpaired *t*-tests with Welch's Correction.

**Supplemental Table 1: Antibodies used in the study**

| Target | Clone | Conjugate | Catalog # | Vendor | Method | Dilution |
|---|---|---|---|---|---|---|
| human CD3 | SK7 | Alexa Fluor 700 | 344822 | Biolegend | Flow Cytometry | 1/50 |
| human CD25 | s20019D | APC | 3385206 | Biolegend | Flow Cytometry | 1/50 |
| human CD45RO | UCHL1 | APC-Cy/7 | 304228 | Biolegend | Flow Cytometry | 1/50 |
| human CD8 | SK1 | BV605 | 344742 | Biolegend | Flow Cytometry | 1/50 |
| human CD45RO | SK3 | BV786 | 344642 | Biolegend | Flow Cytometry | 1/50 |
| human CD25 | BC96 | BV421 | 302630 | Biolegend | Flow Cytometry | 1/50 |
| human CD69 | FN50 | PE | 310906 | Biolegend | Flow Cytometry | 1/50 |
| human CD62L | DREG-56 | PE-Cy/7 | 304822 | Biolegend | Flow Cytometry | 1/50 |
| human CD8 | SK1 | Pacific Blue | 344718 | Biolegend | Flow Cytometry | 1/50 |
| human CD8 | SK1 | APC | 980904 | Biolegend | Flow Cytometry | 1/50 |
| human CCR7 | G043H7 | PerCP/Cy5.5 | 353220 | Biolegend | Flow Cytometry | 1/50 |

| | | | | | | |
|---|---|---|---|---|---|---|
| human CCR7 | G043H7 | BV786 | 353230 | Biolegend | Flow Cytometry | 1/50 |
| human CD107α | H4A3 | BV421 | 328626 | Biolegend | Flow Cytometry | 1/50 |
| human TIM3 | F38-2E2 | PE | 345006 | Biolegend | Flow Cytometry | 1/50 |
| human CD62L | DREG-56 | PE/Cy5 | 304808 | Biolegend | Flow Cytometry | 1/50 |
| human CD62L | SK11 | BV711 | 565040 | BD Horizon | Flow Cytometry | 1/50 |
| human/mouse Granzyme B | QA16A02 | PE/Cy7 | 372214 | Biolegend | Flow Cytometry | 1/50 |
| human CD4 | SK3 | Alexa Fluor 700 | 344622 | Biolegend | Flow Cytometry | 1/50 |
| human PD1 | EH12.2H7 | PE-Dazzle | 329940 | Biolegend | Flow Cytometry | 1/50 |
| human Perforin | DG9 | Pacific Blue | 308118 | Biolegend | Flow Cytometry | 1/50 |
| human CD45 | HI30 | BV570 | 304034 | Biolegend | Flow Cytometry | 1/50 |
| human CD4 | RPA-T4 | PE | 300508 | Biolegend | Flow Cytometry | 1/50 |
| human CCR7 | G043H7 | Alexa Fluor 700 | 353244 | Biolegend | Flow Cytometry | 1/50 |
| human LAG3 | 7H2C65 | APC | 369212 | Biologend | Flow Cytometry | 1/50 |
| human TIGIT | A15153G | PE/Cy7 | 372714 | Biolegend | Flow Cytometry | 1/50 |
| human CD45 | HI30 | BV605 | 304042 | Biolegend | Flow Cytometry | 1/50 |
| human TCF1 | 7F11A10 | APC | 655204 | Biolegend | Flow Cytometry | 1/50 |
| human Granzyme A | CB9 | PE | 989202 | Biolegend | Flow Cytometry | 1/50 |
| Human FASL | NOK-1 | BV421 | 306411 | Biolegend | Flow Cytometry | 1/50 |
| human/mouse TOX | NAN448B | R718 | 569622 | BD Biosciences | Flow Cytometry | 1/50 |
| human/mouse TBET | O4-46 | RB705 | 570294 | BD Biosciences | Flow Cytometry | 1/50 |
| anti-SAPK/JNK Rabbit antibody | | n/a | 9252S | Cell Signaling Technologies | Western Blotting | 1/2000 |
| anti-Phospho-SAPK/JNK (Thr183/Tyr185) | G9 | n/a | 9255S | Cell Signaling Technologies | Western Blotting | 1/2000 |
| anti-c-Jun | 60A8 | n/a | 9165S | Cell Signaling Technologies | Western Blotting | 1/2000 |
| anti-Phospho-c-Jun (Ser73) | D47G9 | n/a | 3270S | Cell Signaling Technologies | Western Blotting | 1/2000 |
| anti-Lamin B1 | D4Q4Z | n/a | 12586 | Cell Signaling Technologies | Western Blotting | 1/15000 |
| Goat anti-Mouse IgG Fc Cross-Adsorbed Secondary Antibody | | HRP | 31439 | Invitrogen | Western Blotting | 1/30000 |
| anti-Rabbit IgG | | HRP | HAF008 | R&D Systems | Western Blotting | 1/15000 |

**Supplemental Table 2: List of parameters and estimation methods for Ordinary Differential Equations model (see: Fig.7)**

| Parameter | Definition | Value | Method of estimation |
|---|---|---|---|
| Tu | Tumor size ($mm^3$) | Day 0 = 20 | 1 $mm^3$ = 100,000 cells (assumption) |
| r | Tumor growth rate | 0.06575 | Fit to untreated tumor curve |
| $Tu_K$ | Carrying capacity of Tumor size | 2000 | Humane endpoint |
| CYT | Cytotoxicity (Tumor cells killed/CAR-T cell/day) | varies | Determined empirically *in vitro* |
| $CYT_d$ | Decay rate of cytotoxicity. Representing exhaustion | -0.876 | Fit to shJNK treated tumor curve |
| CIR | Circulating CAR-T cells | varies | Day 0 = number of CAR-T cells injected |
| $CIR_d$ | Decay rate of CIR population, representing loss of CAR-T cells in circulation | -0.23 | Day 10 ≈ 10% of input. Based on preliminary data |
| a | Infiltration rate of circulating CAR-T cells to the tumor | 0.00035 | Fit to shJNK treated tumor curve |
| TIL | Number of CAR-T cells in the tumor | Day 0 = 0 | n/a. generated based on other parameters. |
| $TIL_g$ | Growth rate of tumor infiltrated CAR-T cells. Representing proliferation in the tumor site | 0.0569 | Fit to shJNK treated tumor curve |
| $TIL_K$ | Carrying capacity of TIL. To limit unrealistic expansion of TILs | 500,000 | assumption |
